# World’s Fastest Brain-Computer Interface: Combining EEG2Code with Deep Learning

**DOI:** 10.1101/546986

**Authors:** Sebastian Nagel, Martin Spüler

**Author notes:** Corresponding author *Email address:* (Sebastian Nagel).

## Abstract

In this paper, we present a Brain-Computer Interface (BCI) that is able to reach an information transfer rate (ITR) of more than 1200 bit/min using non-invasively recorded EEG signals. By combining the EEG2Code method with deep learning, we present an extremely powerful approach for decoding visual information from EEG. This approach can either be used in a passive BCI setting to predict properties of a visual stimulus the person is viewing, or it can be used to actively control a BCI spelling application. The presented approach was tested in both scenarios and achieved an average ITR of 701 bit/min in the passive BCI approach with the best subject achieving an online ITR of 1237 bit/min. The presented BCI is more than three times faster than the previously fastest BCI and allows to discriminate 500,000 different visual stimuli based on 2 seconds of EEG data with an accuracy of up to 100 %. When using the approach in an asynchronous BCI for spelling, we achieved an average utility rate of 175 bit/min, which corresponds to an average of 35 error-free letters per minute. As we observe a ceiling effect where more powerful approaches for brain signal decoding do not translate into better BCI control anymore, we discuss if BCI research has reached a point where the performance of non-invasive BCI control cannot be substantially improved anymore.

## 1. Introduction

A brain-computer interface (BCI) is a device that translates brain signals into output signals of a computer system. The BCI output is mainly used to restore several functionalities of motor disabled people, e.g. for prosthesis control or for communication [1]. Beside the use of BCIs that give the user the ability to actively control a device, passive BCIs have been accepted as a different kind of BCIs that do not have the purpose of voluntary control [2].

In the area of BCIs for communication purposes, BCIs based on visual evoked potentials (VEPs) have emerged as the fastest and most robust approach for BCI communication. Although the idea to use VEPs for BCI control dates back to Vidal [3], Sutter [4] suggested the use of VEPs for a BCI-controlled keyboard in 1984 and showed in 1992 that an ALS patient can use such a system at home to write up to 12 words per minute [5].

Since then, different approaches were demonstrated that use VEPs for BCI control. The majority of VEP-based BCI systems is based on frequency-modulated SSVEPs. The highest information transfer rate (ITR) for an SSVEP-based system was reported by Chen et al. [6], with 267 bit/min on average and up to 319 bit/min for the best subject, which also happens to be the highest ITR being reported for any BCI system, so far.

Instead of using a frequency-modulated visual stimulus for SSVEPs, visual stimuli can also be modulated with a certain pattern to evoke so called code-modulated visual evoke potentials (c-VEPs) that can be used for BCI control. Although Sutter was the first to use this approach [5], it was ignored by BCI research for a long time, and reemerged in 2011 when Bin et al. [7] presented a c-VEP based BCI-system reaching an average ITR of 108 bit/min up to 123 bit/min for the best subject, which was the highest reported ITR at that time. In 2012, Spüler et al. [8] improved the methods for detection of c-VEPs and reached an average ITR of 144 bit/min, up to 156 bit/min for the best subject, which was the highest reported ITR at that time.

A completely different approach to utilize VEPs for BCI control was presented by Thielen et al. [10] in 2015, who created a model to predict the response to short and long visual stimulation pulses and showed that it can be used for BCI control reaching an average ITR of 48 bit/min.

While the model by Thielen et al. was only able to predict responses consisting of short and long pulses, we presented the EEG2Code method [9] to create a model that allows to predict the EEG response to arbitrary stimulation patterns (codes). Using the EEG2Code model for BCI communication resulted in an average ITR of 108 bit/min. In a following publication, we further improved the method and presented an asynchronous BCI with robust non-control state detection to reach an average ITR of 122 bit/min, up to 205 bit/min for the best subject [11].

The EEG2Code method was based on the idea that the VEP response to a complex stimulus is generated by a linear superposition of single-flash VEP responses [12, 13]. In our previous work, we found that we can use a linear method to model the VEP response, but that the VEP response also has non-linear properties that cannot be described by a superposition and cannot be reconstructed with a linear model [9].

As deep learning has become popular in the last years as a powerful machine learning method to create nonlinear prediction models, that outperform other methods in tasks like image classification or speech recognition [14], deep learning methods seem like a natural fit to apply them for neural data.

One class of neural network used in deep learning are convolutional neural networks (CNNs), which were already used in the field of BCIs. The first work which explored CNNs for a BCI is by Cecotti et al. [15], who applied CNNs to P300 data and found CNNs to outperform other methods. In 2017, Kwak et al. [16] proposed a 3-layer CNN that uses frequency features as input for robust SSVEP detection. They compared their CNN approach to other state-of-the-art methods for SSVEP decoding and found CNNs to outperform all of them. Especially for noisy EEG data obtained by a moving participant they achieved an accuracy of 94.03 % compared to 84.65 % achieved by the best compared method. Thomas et al. [17] performed a similar comparison, with the result that the CNN outperformed all other state-of-the-art methods as well.

As we found that the VEP response cannot be appropriately modelled by linear methods, this paper presents an approach to combine the EEG2Code method with deep learning to create a non-linear model that predicts the VEP response to arbitrary stimuli and we show how this method can be used in a BCI.

## 2. Methods

### 2.1. EEG2Code model

In a previous work [9] we proposed a new stimulation paradigm, based on fully random visual stimulation patterns for the use in BCI. In the same work we proposed the EEG2Code model which allows to predicted the stimulation pattern based on the EEG. Furthermore, we have shown how the model can be used for synchronous BCI control. In a subsequent work [11] we have shown that the EEG2Code model can also be used for high-speed asynchronous BCI control. In those works, the EEG2Code model was based on a linear ridge regression. For the sake of completeness, the central parts of the EEG2Code method are explained briefly in the following paragraphs, but a more detailed description can be found in our previous works [11, 9].

The general setup of the BCI is shown in Fig. 1. The EEG2Code method is based on a bit-representation (code) of the visual stimulation pattern, where the properties of the visual stimulus are encoded as one bit (0: black, 1: white). The idea of the EEG2Code method is to predict each bit of the stimulation pattern based on the following 250 ms of EEG data.

**Figure 1:**
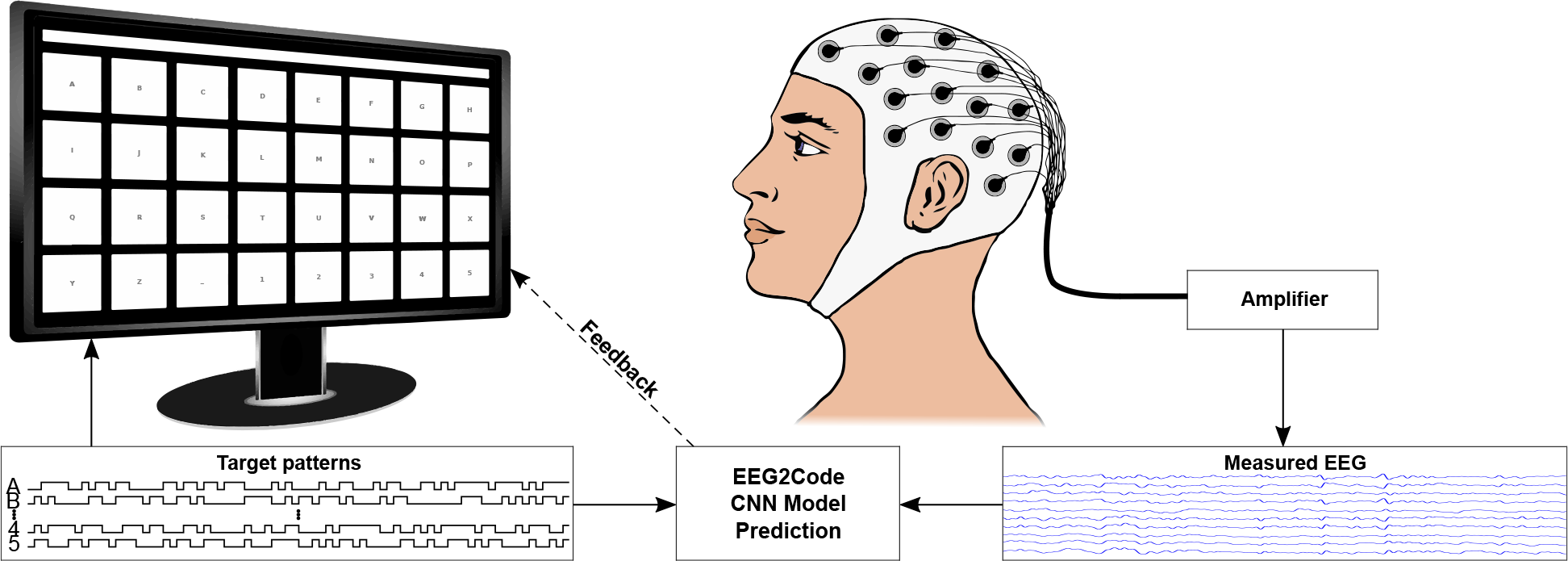
General setup of the BCI speller experiment. The matrix-keyboard layout is as shown on the monitor, it has 32 targets labeled alphabetically from A to Z followed by ‘_’ and numbers 1 to 5. The targets are separated by a blank black space and above targets is the text field showing the written text. Each target is modulated with its own random stimulation pattern. During a trial, the participant has to focus a target. The measured EEG is amplified and afterwards the EEG2Code model predicts the stimulation pattern as shown in Fig. 2. The picture modified from [9]

In our previous works we trained a linear ridge regression method for that purpose, but any regression method can be used. For training the model, the subject has to watch a visual stimulation that is randomly flickering between black and white. After training the EEG2Code model using the bit-sequence of the stimulation pattern and the concurrently recorded EEG signal, the resulting model can be used to predict the property of the visual stimulus in real time. This procedure is also depicted in Fig. 2. In our work we recorded EEG with a sampling rate of 600 Hz and shifted the 250 ms prediction window sample-wise to obtain 600 predictions per second. As the visual stimulus is modulated with 60 Hz (refresh rate of the monitor), we obtained 10 prediction samples for each bit in the stimulation sequence. Those 10 real-valued predictions are averaged to obtain one prediction value for each bit. For the final prediction of the pattern, a threshold of 0.5 is used to either predict the stimulus property as black (0) or white (1).

**Figure 2:**
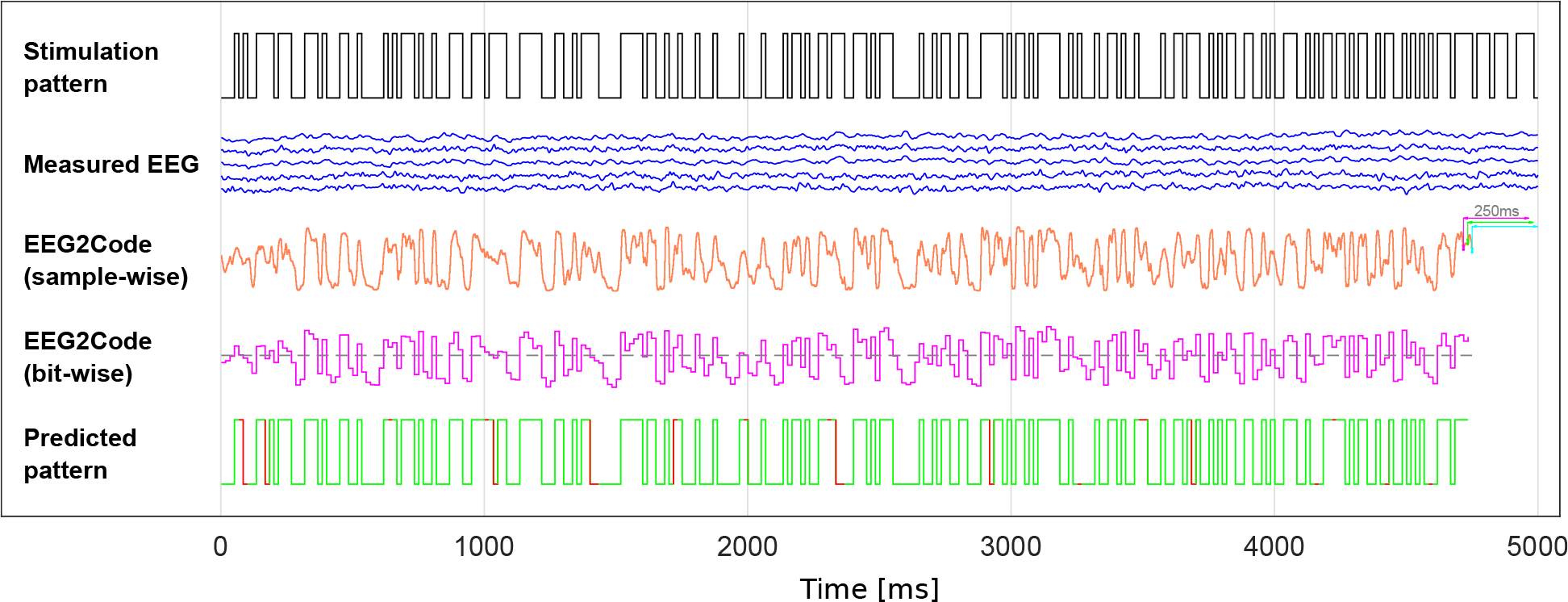
Example of one run where the EEG2Code model with a convolutional neural network (CNN) was used in the passive BCI setting to predicted the visual stimulation pattern. The visual stimulus was modulated with a random stimulation pattern (black line). For each 250 ms window (slided sample-wise) of the measured EEG signals (blue lines), the EEG2Code CNN model predicts a probability value (orange line), which indicates whether the stimulus is black or white. This procedure is shown for 3 exemplary windows (magenta, green, cyan). Note that the model prediction is delayed by 250 ms due to the sliding window approach. The resulting model prediction is now down-sampled to the corresponding number of bits (magenta line). Using a threshold of 0.5 (gray dotted line), the EEG2Code prediction is transformed to the predicted stimulation pattern, which in turn can be compared to the real stimulation pattern (green = match, red = mismatch). For this plot, EEG data and results from the best run were used where an accuracy of 92.6 % was achieved, corresponding to an ITR of 2122 bit/min. An animated version of this figure can be found in the supplementary materials.

The EEG2Code method can either be used in a passive BCI approach to predict the property of the visual stimulus a subject is currently watching (as described above) or it can be used for an active BCI, e.g. a spelling application.

In the BCI control scenario, multiple stimuli (targets) are presented to the subject, where each stimulus is corresponding to a different action (or letter). By comparing the prediction of the EEG2Code model with the stimulation patterns of all targets, we can identify the target that is attended by the subject. For synchronous BCI control, we calculated the correlation between the predicted stimulation pattern and the patterns of all targets, and the target with the highest correlation is selected. For asynchronous BCI control, the EEG2Code prediction is compared to all stimulation patterns continuously. Instead of using the correlation, we calculate the p-values under the hypothesis that the correlation is greater than zero, as this takes the length of a trial into account. If a certain threshold is exceeded, the corresponding target is selected. A more detailed description of the asynchronous classification method can be found in our previous work [11].

### 2.2. Combining EEG2Code with deep learning

Instead of a linear ridge regression that was used in our previous works, we used a convolutional neural network (CNN) as base for the EEG2Code method. The topology of the CNN model is modified from the topology of Gunsteren [18], where it was used for classification of P300 data. Contrary to the EEG2Code ridge regression model, the data is not spatially filtered beforehand. This means the model takes windows of size *T* × *C*, whereby *T* = 150 corresponds to the number of samples (250 ms) and *C* = 32 corresponds to the number of EEG channels. The model consists of five layers, whereby the first layer uses convolutional kernels of size 1 × 32, which means they move over the channels and act as spatial filters. As 16 `Conv` operation are performed, this corresponds to 16 spatial filters in total. The second layer consists of 8 convolutional kernels with a size of 64 × 1 which act as different temporal filters. The remaining layers are used for predicting the stimulation property based on the spatially, and temporally filtered data.

A detailed structure of the EEG2Code CNN is depicted in Fig. 3. It shows each performed operation including the used parameters as well as the input and output dimensions of each operation.

**Figure 3:**
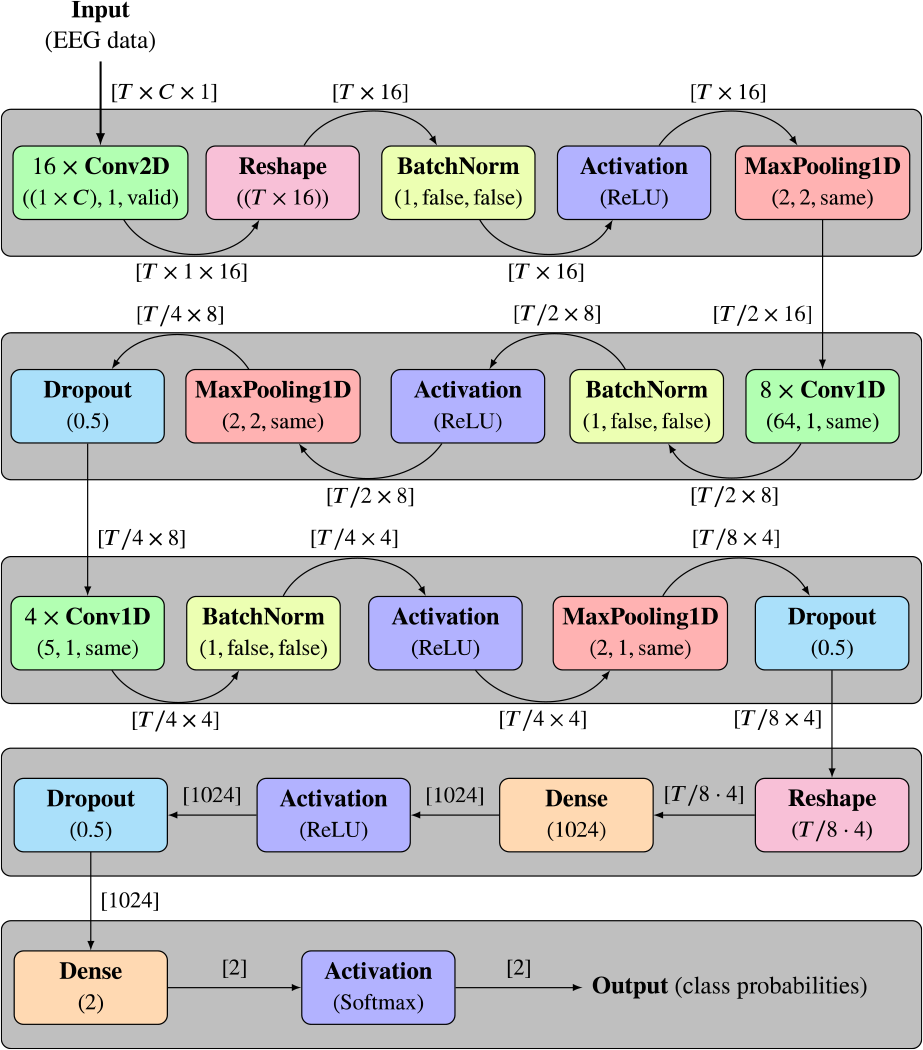
Topology of the EEG2Code convolutional neural network. The five layers are indicated by the gray boxes. Each layer consists of different operations, whereby the same operations are colored the same. Below each operation, the used parameters are given, whereby the parameters for the `Conv` and `MaxPooling` are (`kernel/pooling size, stride, padding`), and for `BatchNorm` (`axis, scale, center`). The edges are labeled with the shape of the input and output data, respectively. The input of the model is a 250 ms (*T* = 150 samples) window of EEG data with *C* = 32 channels. The output are two values which are the probabilities that the input belongs to each class (binary 1 or 0). As the window is shifted sample-wise over the complete trial data, the output is the prediction of the corresponding stimulation pattern.

`MaxPooling` operations are performed to reduce the size of intermediate representation between layers. The `BatchNorm` operation results in a faster and more stable training. The `Dropout` operation is used for regularization to avoid overfitting of the model. The `Dense` operation fully connects the input neurons with the given number of output neurons. The `Activation` operations defines/transforms the output of each neuron depending on a given activation function, whereby the *softmax* function takes an un-normalized vector, and normalizes it into a probability distribution, and therefore gives the probabilities that the initial input belongs to one of the classes. Since the model is trained on two classes (0 or 1), the output are two probabilities *p*_0_ and *p*_1_, one for each class. It must be noted that *p*_1_ = 1 − *p*_0_. The model is trained using a learning rate of 0.001 and a batch size of 256. In total, 25 epochs were trained and the best model was selected as the final model. In this case, the best model refers to the model with the highest validation accuracy, where the validation dataset is independent from the testing dataset. Since the model is trained on two classes (0 or 1), the output are two values *p*_0_ and *p*_1_, which are the probabilities that the input belongs to the corresponding classes, with *p*_1_ = 1 − *p*_0_.

### 2.3. Offline analysis

To evaluate the combination of EEG2Code and deep learning, we evaluated the method offline on different data that was obtained in two previous studies, where the EEG2Code method was uses for synchronous BCI control [9] and an improved version of EEG2Code was demonstrated with asynchronous BCI control [11]. For the offline evaluation, we simulated an online experiment, where the data for training and testing the model was the same as in the online experiment, when the data was recorded.

The data was used for two different scenarios: the active BCI scenario where a target/letter is selected. And the passive BCI scenario, where each bit of the stimulation sequence is predicted. In the latter scenario, the terminology is different and one trial in the passive BCI scenario refers to predicting one bit (16.6 ms stimulus presentation) based on 250 ms of EEG data.

For the synchronous mode, we used a presentation layout with 32 targets arranged as a matrix-keyboard. In total, 384 s of training data were recorded, which means 384 s · 60 bit/s = 23040 random bits were presented. The testing phase was split into 14 runs with a trial duration of 2 s each. Those runs were alternated using fully random stimulation patterns and optimized stimulation patterns, see [9] for details. For the bit prediction accuracy the 7 runs with fully random stimulation were used, whereby for the synchronous BCI control, the 7 runs with optimized stimulation were used. The participants had to perform each run in lexicographic order, with 32 trials per run, so that a total of 448 letters were selected. As we found in our previous study [9] that a trial duration of 1 s is optimal for the synchronous scenario, the simulated online experiment in this study was performed with a trial duration of 1 s.

We further tested how well the EEG2Code method can discriminate 500,000 different stimuli. As each target is modulated with a random stimulation pattern, there is no need to record data from 500,000 targets. Instead, we compared the predicted stimulation pattern, to a set of 500,000 patterns, where one pattern is identical to the one used for stimulation, while the other 499,999 are different random stimulation patterns. Such an analysis was already performed for the EEG2Code ridge regression model in our previous publication [9], where we tested different numbers of stimuli. Although one could test for larger numbers, we stopped at 500,000 due to the increasing computational demands and therefore only used this number for this publication.

For the asynchronous BCI control, the data from [11] was used. The training data also consists of 384 s of recorded EEG data, whereby the optimized stimulation patterns were used. The testing phase consists of 6 runs with 32 trials each and with optimized stimulation patterns. Note that the trial duration varies due to the asynchronous approach, whereby the inter-trial time was set to 500 ms. The trials were performed in lexicographic order. For comparison reasons, the same p-value thresholds were used as determined during the online experiment, see [11] for details.

The experimental setup for recording these data was very similar to the setup described later in this paper, but is also described in more detail in the corresponding publications [11, 9].

### 2.4. Online experiment

To demonstrate that the EEG2Code model using a CNN can also be used in an online BCI to provide real-time feedback, we performed an online experiment. This online experiment should only serve as a proof-of-concept and only one subject was tested.

The subject was the best-performing subject (S01) from our first study. The participation in our first study and the proof-of-concept experiment were approximately 14 month apart.

For training the EEG2Code CNN model, the participant first performed a training phase consisting of 96 runs with 4 s of random stimulation each. Afterwards, 96 runs were performed, whereby each run consisted of 285 trials, with each trial corresponding to one bit of the random stimulation sequence. At the end of the run, an additional 250 ms of random stimulation followed, so that one run had a total length of 5 seconds.

### 2.5. Hardware & Software

The setup was similar to the one used in a previous study [11], except that the EEG2Code CNN computations were performed on an IBM Power System S822LC with four Nvidia^®^ Tesla P100 GPUs using Python v2.7 [19] and the Keras framework [20].

The system consists of a g.USBamp (g.tec, Austria) EEG amplifier, three personal computers (PCs), Brainproducts Acticap system with 32 channels and a LCD monitor (BenQ XL2430-B) for stimuli presentation. Participants are seated approximately 80 cm in front of the monitor.

PC1 is used for the presentation on the LCD monitor, which is set to refresh rate of 60 Hz and its native resolution of 1920 × 1080 pixels. A stimulus can either be black or white, which can be represented by 0 or 1 in a binary sequence and is synchronized with the refresh rate of the LCD monitor, which means each bit of the stimulation patterns are presented for 1/60 s. The timings of the monitor refresh cycles are synchronized with the EEG amplifier by using the parallel port.

As correct synchronisation of EEG and visual stimulation is crucial, we corrected for the monitor raster latency as described in our previous work [21].

PC2 is used for data acquisition, whereby BCI2000 [22] is used as a general framework for recording the data of the EEG amplifier. The amplifier sampling rate was set to 600 Hz, resulting in 10 samples per frame/stimulus. A TCP network connection was established to PC1 in order to send instructions to the presentation layer and to get the modulation patterns of the presented stimuli. During the online experiment, the EEG data was continuously sent to PC3 using a TCP connection. PC3 performs the EEG2Code prediction and sent back the prediction to PC2.

A 32 electrodes EEG layout was used, 30 electrodes were located at Fz, T7, C3, Cz, C4, T8, CP3, CPz, CP4, P5, P3, P1, Pz, P2, P4, P6, PO9, PO7, PO3, POz, PO4, PO8, PO10, O1, POO1, POO2, O2, OI1h, OI2h, and Iz. The remaining two electrodes were used for electrooculography (EOG), one between the eyes and one left of the left eye. The ground electrode (GND) was positioned at FCz and reference electrode (REF) at OZ.

### 2.6. Performance evaluation

The BCI control performance is evaluated using the accuracy, the number of correct letters per minute (CLM) [23], the utility bitrate (UTR) [24] and the information transfer rate (ITR) [25]. The CLM, which take into account that erroneous letters have to be deleted using a backspace symbol, can be computed with the following equation:

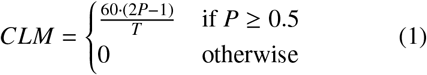

The ITR can be computed with the following equation:

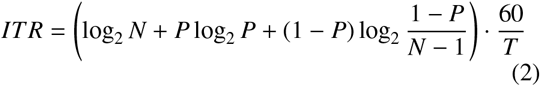

with *N* the number of classes, *P* the accuracy, and *T* the time in seconds required for one prediction. The ITR is given in bits per minute (bit/min). For asynchronous BCI control, *N* equals the number of targets (depending on the layout) and *T* the average trial duration including the inter-trial time. The ITR is generally used to assess how much information a user can convey by BCI control and therefore mostly used as performance measure for BCI communication. However, the ITR also measures how much information can be extracted from the brain signals, which is applicable to all BCI approaches, including passive BCIs.

It should also be noted that the ITR is based on Shannon-Weaver’s model for communication [26], which consists of a information source sending (binary) information that is being encoded, transmitted via a noisy channel, decoded and received by a receiver. As the passive BCI scenario in this paper shows a clearer analogy to Shannon-Weaver’s model than other BCIs, it should be pointed out: Computer A is the information source that decodes binary information in a visual stimulus. The user’s nervous system (eye and brain) is the noisy channel. The BCI acts as a decoder, which decodes the brain signals and the decoded information is received by computer B.

## 3. Results

### 3.1. Offline analysis: simulated online experiment

In the simulated online experiment, we first analyzed the EEG2Code stimulation pattern prediction, which resulted in an average accuracy of 74.9 % using the fully random stimulation patterns and corresponds to an average ITR of 701.3 bit/min. It is worth noting that for S01 an average accuracy of 83.4 % was achieved, which corresponds to 1262.1 bit/min. Additionally, the synchronous BCI control was simulated, which results in average accuracy of 95.9 % using the optimized stimulation patterns and a trial duration of 1 s, which corresponds to an ITR of 183.1 bit/min including the intertrial time of 0.5 s. Detailed results for each subject are listed in Table 1.

**Table 1:** Simulated online results of the EEG2Code method in a passive BCI scenario and for active BCI control. Shown are the average results of all subjects, whereby best results are in bold font. The left part shows the results for the EEG2Code pattern prediction, whereas the right part shows the results for the simulated synchronous BCI control with a trial duration of 1 s. For both, the previous results using the ridge regression model as well as the new results using the CNN model are shown. For all, the accuracies (ACC) and the corresponding ITRs are given. The ITRs are calculated using Eq. 2 with *N* = 2 (*N* = 32) and *T* = 1/60*s* (*T* = 1.5*s*).

| Subject | Pattern prediction |  |  |  | Synchronous BCI control |  |  |  |
| --- | --- | --- | --- | --- | --- | --- | --- | --- |
|  | Ridge regression |  | CNN |  | Ridge regression |  | CNN |  |
|  | ACC [%] | ITR [bpm] | ACC [%] | ITR [bpm] | ACC [%] | ITR [bpm] | ACC [%] | ITR [bpm] |
| S01 | **69.1** | **389.9** | **83.4** | **1262.1** | **99.2** | **195.8** | **100.0** | **200.0** |
| S02 | 64.5 | 222.4 | 72.9 | 567.3 | 94.1 | 175.5 | 94.6 | 177.3 |
| S03 | 63.7 | 196.5 | 72.5 | 545.7 | 83.4 | 141.2 | 95.5 | 180.6 |
| S04 | 65.6 | 257.8 | 76.8 | 787.9 | 95.9 | 182.1 | 98.7 | 193.2 |
| S05 | 66.3 | 282.7 | 78.7 | 908.9 | 96.4 | 183.8 | 99.6 | 197.5 |
| S06 | 67.1 | 308.2 | 75.4 | 704.9 | 98.1 | 191.0 | 98.7 | 193.2 |
| S07 | 63.7 | 196.8 | 68.5 | 363.9 | 88.4 | 156.4 | 87.9 | 154.9 |
| S08 | 60.9 | 124.5 | 77.1 | 807.1 | 65.6 | 94.6 | 99.6 | 197.5 |
| S09 | 60.2 | 109.4 | 68.5 | 363.7 | 53.5 | 68.0 | 87.5 | 153.5 |
| mean | 64.6 | 232.0 | 74.9 | 701.3 | 86.1 | 154.3 | 95.9 | 183.1 |

Furthermore, also the asynchronous BCI control was simulated using the 32-target matrix-keyboard layout. The results for each participant are shown in Table 2. The average target prediction accuracy is 98.5 % (ITR: 175.5 bit/min) with an average trial duration of 1.71 s (including 0.5 s inter-trial time). In total, 91.6 % of all trials could be classified faster compared to the ridge regression model. Finally, this results in an average spelling speed of 35.3 correct letters per minute (CLM) with a maximum of 48.2 CLM, which corresponds to a utility rate of 175 bit/min and 239 bit/min, respectively.

**Table 2:** Simulated online results for an asynchronous BCI speller. Shown are the results for the lexicographic-spelling (matrix-layout, 32 targets). The number of correct letters per minute (CLM), the target prediction accuracy, the information transfer rates (ITR), the average trial duration (including an inter trial time of 0.5 s) and the utility bitrate (UTR). Best results are in bold font.

| Subject | CLM | Accuracy [%] | ITR [bpm] | Time [s] | UTR [bpm] |
| --- | --- | --- | --- | --- | --- |
| S10 | 33.3 | 99.0 | 165.4 | 1.77 | 165.0 |
| S11 | **48.2** | 97.9 | **238.9** | **1.19** | **238.8** |
| S12 | 39.7 | 99.5 | 197.8 | 1.49 | 196.8 |
| S13 | 28.7 | 99.5 | 142.8 | 2.07 | 142.1 |
| S14 | 35.8 | 95.3 | 177.8 | 1.52 | 177.6 |
| S15 | 30.7 | 99.0 | 152.4 | 1.92 | 152.0 |
| S16 | 45.1 | **100.0** | 225.3 | 1.33 | 223.2 |
| S17 | 25.8 | 97.4 | 127.8 | 2.21 | 127.7 |
| S18 | 30.9 | **100.0** | 154.4 | 1.94 | 153.0 |
| S19 | 34.8 | 97.4 | 172.3 | 1.64 | 172.2 |
| mean | 35.3 | 98.5 | 175.5 | 1.71 | 174.8 |

### 3.2. Discriminating 500,000 different stimuli

The results for using the EEG2Code approach for the discrimination of 500,000 different stimuli patterns can be seen in Table 3. While the EEG2Code method with ridge regression was able to identify the correct stimulus with an average accuracy of 54.9 %, the EEG2Code deep learning approach achieved an average accuracy of 84.0 %. Notably, subject S01 achieved an accuracy of 100 % showing that for all trials, the correct stimulus pattern was identified.

**Table 3:**
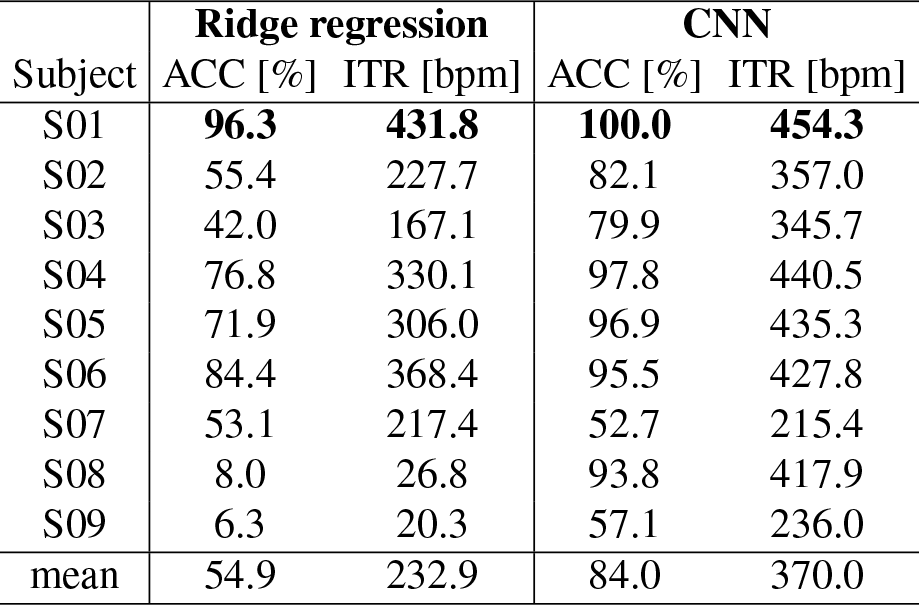
Simulated online results of the EEG2Code method for the discrimination of 500,000 different stimuli. Shown are the average results of all subjects, whereby best results are in bold font. The results are for the simulated synchronous BCI control with 500,000 simulated targets based on 2 s of EEG data. The accuracies (ACC) and the corresponding ITRs are given, whereby the ITRs are calculated using Eq. 2 with *N* = 500000 and *T* = 2.5*s*.

| Subject | Ridge regression |  | CNN |  |
| --- | --- | --- | --- | --- |
|  | ACC [%] | ITR [bpm] | ACC [%] | ITR [bpm] |
| S01 | **96.3** | **431.8** | **100.0** | **454.3** |
| S02 | 55.4 | 227.7 | 82.1 | 357.0 |
| S03 | 42.0 | 167.1 | 79.9 | 345.7 |
| S04 | 76.8 | 330.1 | 97.8 | 440.5 |
| S05 | 71.9 | 306.0 | 96.9 | 435.3 |
| S06 | 84.4 | 368.4 | 95.5 | 427.8 |
| S07 | 53.1 | 217.4 | 52.7 | 215.4 |
| S08 | 8.0 | 26.8 | 93.8 | 417.9 |
| S09 | 6.3 | 20.3 | 57.1 | 236.0 |
| mean | 54.9 | 232.9 | 84.0 | 370.0 |

### 3.3. Online experiment

To demonstrate that the EEG2Code CNN model can also be used in an online BCI, we performed an experiment where we invited the best subject from the first experiment to participate. Averaged over all runs, an average bit prediction accuracy of 83.4 % was achieved which corresponds to an average ITR of 1237 bit/min using *N* = 2 and *T* = 5/285 s, taking into account that 285 trials are predicted in a 5 s run. The distribution of the average performance per run are depicted in Fig. 4. The results also shows that the prediction of the run with the best performance had an accuracy of 92.6 % and an ITR of 2122 bit/min.

**Figure 4:**
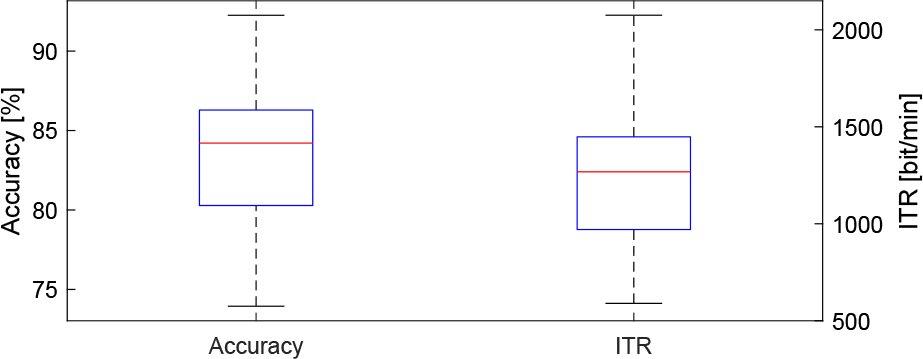
Bit prediction accuracies and corresponding ITRs achieved during an online experiment of subject S01. Shown is the distribution of the average performance per run with the red line representing the median. The data consists of 96 runs, with each running having 285 trials (bits). The ITRs were calculated using Eq. 2 with *N* = 2 and *T* = 5/285 *s*.

## 4. Discussion

In this work, we have combined the EEG2Code method with deep learning to show that this approach can be used in two different kinds of BCI: a passive BCI to predict properties of the visual stimulus the user is viewing, or for controlling a BCI to select letters.

In an online experiment, we were able to show that the approach is online capable and that a subject can reach 1237 bit/min when using the method in a passive BCI.

In an offline analysis, we compared the EEG2Code method using a ridge regression against the EEG2Code method with deep learning and could show an increase in the classification accuracy from 64.6 % to 74.9 % for the pattern prediction. With regard to the information that can be extracted from the EEG, the ITR could be improved by 202 % with deep learning (from 232 bit/min to 701 bit/min).

Compared to current state-of-the-art approaches, the EEG2Code deep learning approach clearly outperforms the previously fastest system by Chen et al. [6]. They reported the previously highest ITR for a BCI, with an average ITR of 267 bit/min and an online ITR of 319 bit/min for the best subject, which we could raise to 701 bit/min and 1237 bit/min, respectively.

### 4.1. Ceiling effect

When comparing the results of the passive BCI for pattern prediction (701 bit/min) with the results for BCI control (183 bit/min), we observe a ceiling effect. Although we are able to extract an average of 701 bit/min of information from the EEG, we can only use 183 bit/min of this information for BCI control. The discrepancy between extracted information and BCI control becomes even larger when looking at single subjects. The worst subject (S09) had an ITR of 364 bit/min for the pattern prediction, which translates into an ITR of 153 bit/min for BCI control. The best subject (S01) had an ITR of 1262 bit/min for the pattern prediction, which translates into an ITR of 200 bit/min. Comparing those two subjects shows that an increase of 898 bit/min for the pattern prediction translates into an increase of only 47 bit/min for BCI control.

Due to the limited number of targets (N=32) and trial duration given by the BCI communication system (T=1.5; 1s trial + 0.5 pause), the ITR is limited to a maximum of 200 bit/min. This limit can only be increased by either reducing the trial duration or by increasing the number of targets. For chasing higher ITRs in the lab, the trial duration could be reduced to 1 s (0.5 s trial + 0.5 s pause), which corresponds to a maximum ITR of 300 bit/min, but with values below 1 s the system becomes too fast to be realistically usable.

Increasing the number of targets is another way to increase the ITR. The EEG2Code method has the unique property that it allows a virtually unlimited amount of targets (e.g. 2^120^ = 1.3 · 10^36^ targets with 2 s of visual stimulation). In this work, we simulated an online BCI with *N* = 500, 000 targets and could show that in this case, the performance of the EEG2Code ridge regression model translates completely to BCI control with 232.0 bit/min for the pattern prediction and 232.9 bit/min for BCI control. But when using deep learning, we again observe a ceiling effect with 701 bit/min for the pattern prediction translated into 370 bit/min for BCI control. While it is likely that the ITR for BCI control would increase further when using more targets, it should be pointed out that this is just a theoretical exercise, because communication systems with such a higher number of targets are practically unusable. Nevertheless, these simulations underscore the power of the EEG2Code method as even with such a large number of targets, high accuracies were reached. In case of the best subject, we could discriminate 500,000 different stimuli with an accuracy of 100 % based on 2 s of EEG data.

While there theoretically are ways to overcome the ceiling effect, real BCI control is still limited by minimum trial duration and a maximum number of targets which are usable. While those limits may vary between users, it is likely that 60 characters per minute is a limit that cannot be surpassed significantly by non-invasive BCI systems (or invasive VEP-based systems).

### 4.2. Brain signal decoding vs. BCI communication

As Sutter has shown in 1992 [5] that an ALS patient can use a VEP-based BCI at home to write up to 12 words (about 60 characters) per minute, cynical voices might argue that BCI research hasn’t made any progress since then as there is still no system that allows higher communication rates. While the limit for VEP-based BCI communication seems to have already been reached in 1992, the methods for decoding brain signals have evolved since then.

Currently, in the BCI literature there is no distinction being made between the performance for brain signal decoding and BCI communication. This wasn’t necessary because all information that could be extracted from the EEG could be used for BCI control so that the performance of a method from a decoding perspective was equal to the performance of a method from the perspective of BCI control. However, this has changed due to the ceiling effect that we observed in this work.

We therefore argue that BCI research should make a clearer distinction between those two perspectives. From the perspective of brain signal decoding, the aim is to decode the signal as good as possible, which means to extract as much information from signals as possible. Therefore, the information transfer rate (ITR) is the perfect measure to evaluate a method in terms of its signal decoding power. When evaluating a method in terms of decoding power, it is also irrelevant if it is being used for BCI control, in a passive BCI or for other purposes.

From a perspective of BCI communication, the ITR is not a good measure for communication performance [24] as it is a pure measure of information that does not take into account how humans use a communication systems (e.g. backspace to correct errors). For this reason, metrics like the utility metric [24] or correct letters per minute [23] are a more appropriate measure. Further, and more importantly, when evaluating a BCI system with regard to its communication performance, it should be ensured that the system can be used in an end-user scenario. From our perspective, end-user scenario is not limited to the use by patients, but we refer to any user. With the term end-user scenario we want to stress the importance of a non-control state detection that detects if a user wants to currently use the BCI or not. In an end-user scenario, there are always time periods where the user does not want to control the BCI either because they are currently listening to another person, reading a webpage or sitting in their wheelchair and enjoying a beautiful sunset. Without a non-control state detection the BCI will output random garbage, click random webpage-links, or drive the wheelchair off in random directions, which is why a non-control state detection is essential for end-user BCIs.

With regard to the work of Chen et al. [6] who presented a BCI to reach an average ITR of 267 bit/min, we don’t see their system as end-user suitable as it does not have a non-control state detection. Further, the user only has 280 ms between letter selection and the start of the next trial which is insufficient to react to errors and is only usable by highly trained subjects. We thereby consider the system by Chen et al. [6] as a pure technical demonstration of a brain signal decoding approach.

In contrast, we have shown the EEG2Code method to be usable in an asynchronous BCI with robust non-control state detection [11] and achieve average communication speeds of 35 correct letters per minute (utility bitrate of 175 bit/min) and thereby consider the BCI be the fastest end-user suitable system for non-invasive BCI communication. For comparison, the previously fastest non-invasive BCI system, that can be considered end-user suitable, was presented by Suefusa et al. [27] and reached an average ITR of 67.7 bit/min.

We encourage fellow researchers to make a clearer distinction between signal decoding performance and BCI control performance and to ensure that the BCI system is viable in an end-user scenario when evaluating BCI control performance. As we have demonstrated a ceiling effect and that it becomes increasingly difficult to improve BCI control performance by better signal decoding, other approaches for improving BCI communication, like language modelling or predictive spelling, gain more importance and further demonstrate the need to separate signal decoding performance from the performance of BCI communication.

### 4.3. Conclusion

In this paper, we have presented a novel approach that combines deep learning with the EEG2Code method to predict properties of a visual stimulus from EEG signals. We could show that a subject can use this approach in an online BCI to reach an information transfer rate (ITR) of 1237 bit/min, which makes the presented BCI system the fastest system by far. In a simulated online experiment with 500,000 targets we could further show that the presented method allows to differentiate 500,000 different stimuli based on 2 s of EEG data with an accuracy of 100 % for the best subject. As the presented method can extract more information from the EEG than can be used for BCI control, we discussed a ceiling effect that shows that more powerful methods for brain signal decoding do not necessarily translate into better BCI control and that it is important to differentiate between the performance of a method for decoding brain signals and its performance for BCI control.

## Data availability

All data used in this paper were made publicly available. The data can be found online separated for the synchronous simulated online experiment [28], the asynchronous simulated online experiment [29], and the online BCI experiment [30].

## Code availability

To demonstrate that our results can be reproduced, we made a python script publicly available [30] that allows to reproduce the results from the online passive BCI experiment. The script predicts the stimulation patterns for each run. The attached CNN model is the one used online. When running the script to train a new CNN model, the results may slightly differ due to random properties of the method (e.g. dropout learning).

## Supporting information

Supplemental Video 1 (animated version of Fig. 2)

## Acknowledgments

This work was supported by the German Research Council (DFG; SP 1533/2-1). The server running the deep-learning framework was provided by the *IBM Shared University Research Grant*.

## References

[1] J. R. Wolpaw, N. Birbaumer, D. J. McFarland, G. Pfurtscheller, T. M. Vaughan, Brain–computer interfaces for communication and control, Clinical neurophysiology 113 (6) (2002) 767–791.

[2] T. O. Zander, C. Kothe, Towards passive brain–computer interfaces: applying brain–computer interface technology to human–machine systems in general, Journal of neural engineering 8 (2) (2011) 025005.

[3] J. J. Vidal, Real-time detection of brain events in eeg, Proceedings of the IEEE 65 (5) (1977) 633–641.

[4] E. E. Sutter, The visual evoked response as a communication channel, in: Proceedings of the IEEE Symposium on Biosensors, 1984, pp. 95–100.

[5] E. E. Sutter, The brain response interface: communication through visually-induced electrical brain responses, Journal of Microcomputer Applications 15 (1) (1992) 31–45.

[6] X. Chen, Y. Wang, M. Nakanishi, X. Gao, T.-P. Jung, S. Gao, High-speed spelling with a noninvasive brain–computer interface, Proceedings of the national academy of sciences 112 (44) (2015) E6058–E6067. doi:10.1073/pnas.1508080112.

[7] G. Bin, X. Gao, Y. Wang, Y. Li, B. Hong, S. Gao, A high-speed bci based on code modulation vep, Journal of neural engineering 8 (2) (2011) 025015.

[8] M. Spüler, W. Rosenstiel, M. Bogdan, Online adaptation of a c-vep brain-computer interface (bci) based on error-related potentials and unsupervised learning, PloS one 7 (12) (2012) e51077.

[9] S. Nagel, M. Spüler, Modelling the brain response to arbitrary visual stimulation patterns for a flexible high-speed brain-computer interface, PloS one 13 (10) (2018) e0206107. doi:10.1371/journal.pone.0206107.

[10] J. Thielen, P. van den Broek, J. Farquhar, P. Desain, Broad-band visually evoked potentials: re (con) volution in brain-computer interfacing, PloS one 10 (7) (2015) e0133797.

[11] S. Nagel, M. Spüler, Asynchronous non-invasive high-speed bci speller with robust non-control state detection, bioRxivdoi:10. 1101/489013.

[12] E. C. Lalor, B. A. Pearlmutter, R. B. Reilly, G. McDarby, J. J. Foxe, The vespa: a method for the rapid estimation of a visual evoked potential, Neuroimage 32 (4) (2006) 1549–1561. doi:10.1016/j.neuroimage.2006.05.054.

[13] A. Capilla, P. Pazo-Alvarez, A. Darriba, P. Campo, J. Gross, Steady-state visual evoked potentials can be explained by temporal superposition of transient event-related responses, PLOS ONE 6 (1) (2011) 1–15. doi:10.1371/journal.pone.0014543.

[14] K. Simonyan, A. Zisserman, Very deep convolutional networks for large-scale image recognition, arXiv preprint arXiv:1409.1556.

[15] H. Cecotti, A. Graser, Convolutional neural networks for p300 detection with application to brain-computer interfaces, IEEE transactions on pattern analysis and machine intelligence 33 (3) (2011) 433–445.

[16] N.-S. Kwak, K.-R. Müller, S.-W. Lee, A convolutional neural network for steady state visual evoked potential classification under ambulatory environment, PloS one 12 (2) (2017) e0172578.

[17] J. Thomas, T. Maszczyk, N. Sinha, T. Kluge, J. Dauwels, Deep learning-based classification for brain-computer interfaces, in: Systems, Man, and Cybernetics (SMC), 2017 IEEE International Conference on, IEEE, 2017, pp. 234–239.

[18] F. van Gunsteren, Deep neural networks for classification of eeg data, Master’s thesis, University of Tübingen, WSI, Tübingen (July 2018).

[19] Python, version 2.7, Python Software Foundation, Wilmington, Delaware, 2010.

[20] F. Chollet, et al., Keras, keras.io (2015).

[21] S. Nagel, W. Dreher, W. Rosenstiel, M. Spüler, The effect of monitor raster latency on veps, erps and brain–computer interface performance, Journal of neuroscience methods 295 (2018) 45–50.

[22] G. Schalk, D. J. McFarland, T. Hinterberger, N. Birbaumer, J. R. Wolpaw, Bci2000: a general-purpose brain-computer interface (bci) system, IEEE Transactions on biomedical engineering 51 (6) (2004) 1034–1043. doi:10.1109/TBME.2004. 827072.

[23] M. Spüler, A high-speed brain-computer interface (bci) using dry eeg electrodes, PloS one 12 (2) (2017) e0172400.

[24] B. Dal Seno, M. Matteucci, L. T. Mainardi, The utility metric: a novel method to assess the overall performance of discrete brain–computer interfaces, IEEE Transactions on Neural Systems and Rehabilitation Engineering 18 (1) (2010) 20–28.

[25] J. R. Wolpaw, H. Ramoser, D. J. McFarland, G. Pfurtscheller, Eeg-based communication: improved accuracy by response verification, IEEE transactions on Rehabilitation Engineering 6 (3) (1998) 326–333. doi:10.1109/86.712231.

[26] C. E. Shannon, A mathematical theory of communication, Bell system technical journal 27 (3) (1948) 379–423.

[27] K. Suefusa, T. Tanaka, Asynchronous brain–computer interfacing based on mixed-coded visual stimuli, IEEE Transactions on Biomedical Engineering 65 (9) (2018) 2119–2129.

[28] S. Nagel, M. Spüler, Modelling the brain response to arbitrary visual stimulation patterns for a flexible high-speed BrainComputer Interface doi:10.6084/m9.figshare.7058900.v1.

[29] S. Nagel, M. Spüler, Asynchronous non-invasive high-speed BCI speller with robust non-control state detection doi:10. 6084/m9.figshare.7611275.v1.

[30] S. Nagel, M. Spüler, World’s Fastest Brain-Computer Interface: Combining EEG2Code with Deep Learning doi:10.6084/m9.figshare.7701065.v1.

